# The spatiotemporal dynamics of recognition memory for complex versus simple auditory sequences

**DOI:** 10.1101/2022.05.15.492038

**Authors:** G. Fernández Rubio, E. Brattico, S. A. Kotz, M. L. Kringelbach, P. Vuust, L. Bonetti

## Abstract

Differently from visual recognition, auditory recognition is a process relying on the organization of single elements that evolve in time. Here, we aimed to discover the spatiotemporal dynamics of this cognitive function by adopting a novel strategy for varying the complexity of musical sequences. We selected traditional tonal musical sequences and altered the distance between pitches to obtain matched atonal sequences. We then recorded the brain activity of 71 participants using magnetoencephalography (MEG) while they listened to and later recognized auditory sequences constructed according to simple (tonal) or complex (atonal) conventions. Results reveal qualitative changes in neural activity dependent on stimulus complexity: recognition of tonal sequences engaged hippocampal and cingulate areas, whereas recognition of atonal sequences mainly activated the auditory processing network. Our findings highlight the involvement of a cortico-subcortical brain network for auditory recognition and support the idea that stimulus complexity qualitatively alters the neural pathways of recognition memory.

## Introduction

Encoding and recognizing sounds that are structurally complex is a cognitive challenge relying on neural mechanisms that are not yet fully elucidated. To memorize complex sound sequences, we likely depend on the temporal organization of a stimulus’ components and memory functions ^1^.

Memory encoding takes place in the hippocampus ^2-4^, whereas subsequent processes related to recognition memory are supported by a functional network of interconnected regions in the medial temporal lobe, including the hippocampus, insula, and inferior temporal cortex ^2, 5, 6^. For memory consolidation, communication between hippocampal and neocortical areas is needed ^7-9^. Much evidence comes from studies using static visual stimuli, such as pictures of objects, faces, or natural scenes ^10-12^. In audition, however, information and meaning unfold over time as the brain attempts to predict upcoming stimuli based on prior memory representations. Hence, to better understand memory recognition and its underlying fast brain dynamics, novel methods must be adopted that highlight the temporal properties of dynamic stimuli. This can be done by studying the neural activity underlying the processing of sound sequences that acquire meaning through their evolution over time, such as music ^13-15^.

According to the predictive coding of music (PCM) theory, music processing is bound by hierarchical Bayesian rules, wherein the brain compares musical information with its internal predictive model in an attempt to reduce a prediction error ^16-19^. Specifically, bottom-up sensations evoked by auditory stimuli are processed in primary cortices and contrasted with top-down predictions in higher-order cortices to generate musical expectations and minimize hierarchical prediction errors ^19-21^. Predictive mechanisms rely on long-and short-term memory functions, familiarity, and listening strategies to create musical expectations ^18^. Overall, the PCM model provides a reliable framework for studying music perception ^22-25^, training ^26, 27^, action ^8, 29^, synchronization ^30-32^, and emotion ^33-35^. In recent years, studies have also began exploring the neural underpinnings of musical memory. Using functional resonance imaging (fMRI) and a naturalistic music listening paradigm, Alluri et al. ^36^ investigated the neural correlates of music processing and reported activation of cognitive, motor, and limbic brain networks for the continuous processing of timbral, tonal, and rhythmic features. Subsequently, using the same stimuli, Burunat et al. ^37^ reported the recruitment of memory-related and motor brain regions during the recognition of musical motifs. Despite their contributions, these studies fail to identify the fine-grained temporal mechanisms of sound encoding and memory processes.

More recently, we introduced novel applications of magnetoencephalography (MEG) combined with magnetic resonance imaging (MRI) to study music recognition. These studies accentuated the temporal involvement of a widespread cortico-subcortical brain network comprising the primary auditory cortex, superior temporal gyrus, frontal operculum, cingulate gyrus, orbitofrontal cortex, and hippocampus during recognition of auditory (musical) sequences ^38-40^. Overall, these investigations have provided unique insight into the neural mechanisms underlying the recognition of temporal sequences. What remains to be addressed is how these mechanisms are modulated by stimulus complexity.

Here, we used melodic sequences, where meaning emerged from the sequential combination of individual tones over time ^40^, and varied the tone distribution to obtain new, complex musical sequences. In this scenario, encoding and recognition of the musical sequences largely depend on the sequential order of the tones that comprise it. We first selected musical sequences based on the rules of tonality, which is the dominant musical system in Western popular music ^41^. Second, by modifying the tone intervals (i.e., the distances between pitches) while keeping all other variables (e.g., rhythm, tempo, timbre) constant, we generated matched *atonal* musical sequences. The stimulus manipulation was based on previous literature, which reported that tonal rather than atonal musical sequences are overall easier to process ^42-46^ and more appreciated by non-expert listeners ^46-48^. Unlike tonal music, atonal music is characterized by the absence of a clear tonal center and hierarchical stability, which significantly reduces its predictive value and gives rise to increased prediction errors ^42-45, 49^. Thus, we expected that the alteration of tonal intervals would reduce the predictability of the atonal sequences, leading to increased difficulty to recognize them.

To summarize, in the current study we used magnetoencephalography (MEG) and a musical recognition task ^38-40^ while participants listened to and recognized auditory (musical) sequences of varying complexity. We aimed at describing its fine-grained spatiotemporal dynamics. Following previous studies ^36-40^, we expected that the recognition of auditory sequences would activate a widespread brain network that includes both auditory (e.g., primary auditory cortex, superior temporal gyrus, Heschl’s gyrus, planum temporale, insula) and memory processing areas (e.g., hippocampus, cingulate gyrus). We further hypothesized that neural activity would be distributed along two main frequency bands that reflected the occurrence of two different cognitive processes: a slow frequency band related to the recognition of the full musical sequence in memory processing areas, and a fast frequency band associated with the processing of each individual tone of the musical sequence in auditory regions. More importantly, we hypothesized that, based on stimulus complexity, tonal music would be more efficiently processed than atonal music, which would be reflected in different behavioral responses and distinct neural pathways during recognition of tonal and atonal sequences.

## Results

### Behavioral data

Participants performed an old/new auditory recognition task. They first listened to a full musical piece (encoding) and subsequently identified which musical sequences were memorized or novel. During recognition, the response accuracy and reaction time of the participants were recorded using a joystick. These behavioral data were statistically analyzed to examine the differences between the four experimental conditions (memorized tonal sequences, novel tonal sequences, memorized atonal sequences, novel atonal sequences).

A one-way analysis of variance (ANOVA) showed that the differences in response accuracy were statistically significant, *F*(3, 280) = 6.87, *p* = .002. Post-hoc analyses indicated that the average number of correct responses was significantly lower for memorized atonal sequences (M = 30.98, SD = 5.46) than for novel atonal (M = 34.51, SD = 4.26, *p* < .001), memorized tonal (M = 34.34, SD = 5.95, *p* = .002) and novel tonal sequences (M = 34.41, SD = 6.04, *p* = .001). **Figure 2A** shows the average number of correct responses, standard deviation, and statistically significant differences per condition.

**Figure 1.**
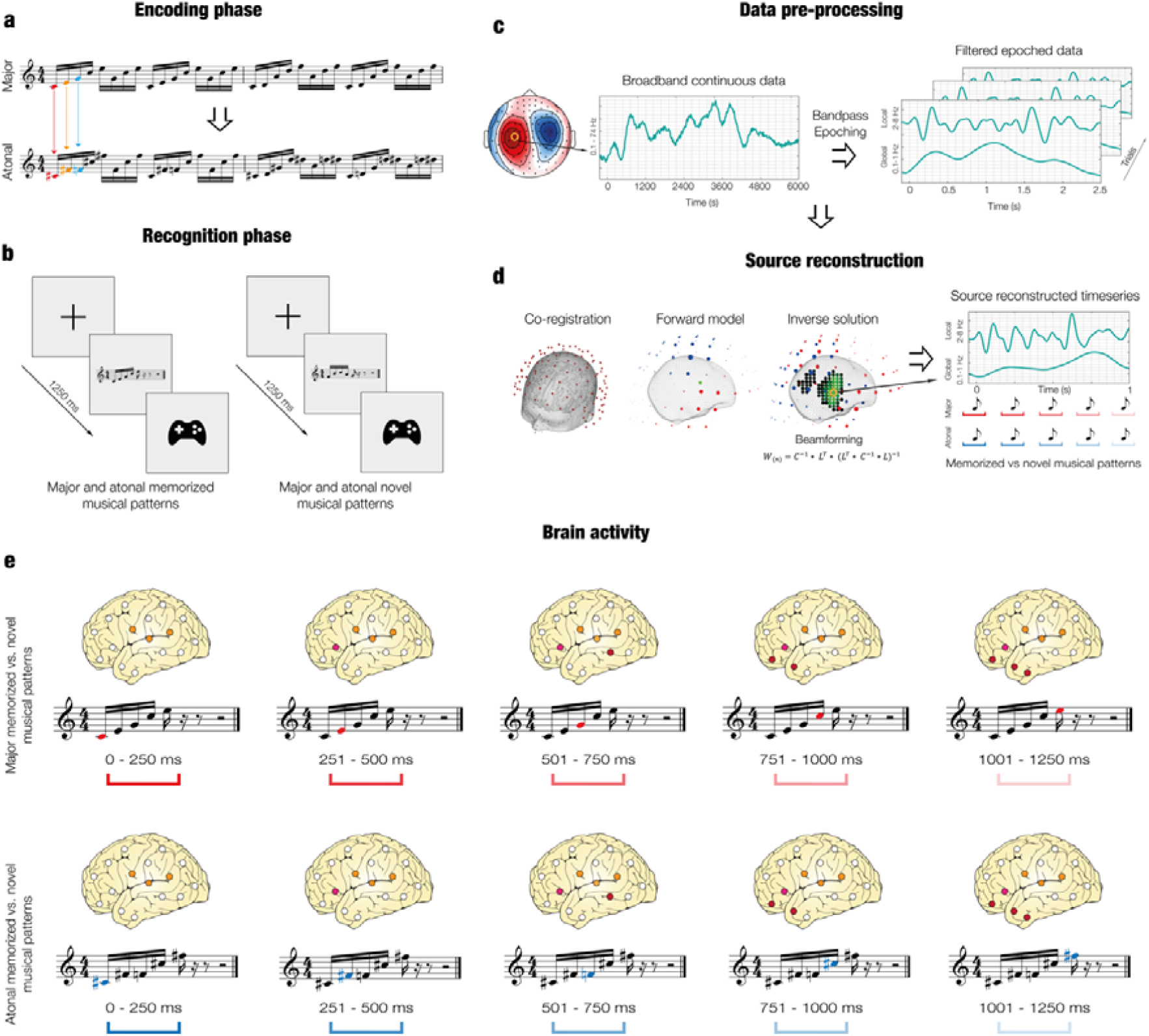
Experimental stimuli and design, data analyses and (temporal) brain activity. **a** – Two musical pieces were used in the experiment: the right-hand part of J. S. Bach’s Prelude No. 1 in C Major, BWV 846 (i.e., “tonal”, top row), and an atonal version of the prelude (i.e., “atonal”, bottom row). Both pieces were matched in terms of the sequential presentation of the tones, rhythmic patterns, dynamics, and duration, and their melodic contour was almost identical. The atonal piece was created by LB by assigning new tones that were one or two semitones lower or higher than the original tones of the tonal piece. For example, C (in red) was converted into C sharp (in red), E (in orange) was converted into F sharp (in orange), G (in blue) was converted into F (in blue), etc. **b** – Participants performed the experimental task twice (once for the tonal piece and once for the atonal piece) and the order of presentation was randomized across participants. After listening to the full piece, participants were presented with excerpts that belonged to the piece or with new excerpts and were asked to state whether the excerpts were “memorized” or “novel” using a joystick. **c** – The task was administered to the participants while their brain activity was recorded using MEG. The continuous neural data was preprocessed. **d** – Source reconstruction analyses were conducted to identify the brain sources that generated the neural activity. The data was first bandpass-filtered into two frequency bands (0.1 – 1 Hz and 2 – 8 Hz) and the MEG and MRI data were co-registered. An overlapping-spheres forward model was computed using an 8-mm grid and a beamforming algorithm was applied as the inverse solution. Finally, the source reconstructed time series was computed for both tonal and atonal data and their contrast in both frequency bands. **e** – Contrasts between memorized and novel sequences were calculated for each tone that comprised the tonal and atonal musical sequences for both frequency bands.

**Figure 2.**
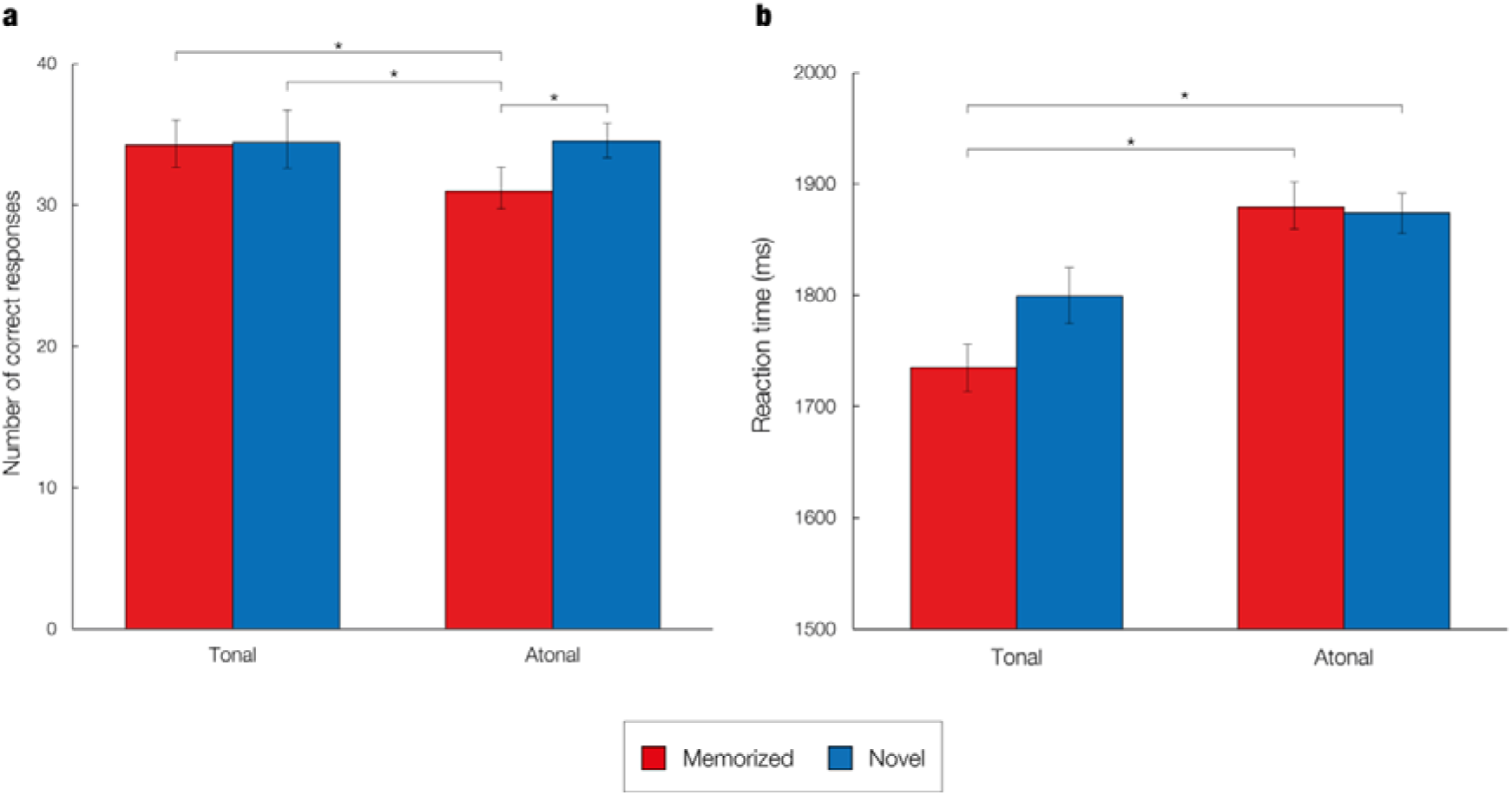
Analyses of behavioral data. **a** – Average number of correct responses for each of the experimental conditions. Asterisks denote a statistically significant difference between two conditions. Error bars show the standard deviation. **b** – Average reaction times for each of the experimental conditions. Asterisks denote a statistically significant difference between two conditions. Error bars show the standard deviation.

Regarding the mean reaction time, there was a statistically significant difference between conditions as determined by one-way ANOVA, *F*(3, 280) = 4.94, *p* = .002. Post-hoc analyses revealed that the average reaction time was significantly lower for memorized tonal sequences (M = 1735.17, SD = 259.91) compared to memorized atonal (M = 1879.44, SD = 259.34, *p* = .005) and novel atonal sequences (M = 1873.78, SD = 250.48, *p* = .007), but not compared to novel tonal sequences (M = 1799.52, SD = 267.14, *p* = .450). **Figure 2B** displays the mean reaction time, standard deviation, and statistically significant differences per condition.

### MEG sensor data

The MEG data (204 planar gradiometers and 102 magnetometers) were analyzed at the MEG sensor level, using the broadband signal. Although the emphasis of the study lays in identifying the brain areas involved in recognizing tonal versus atonal musical sequences, the MEG sensor data were examined to assess whether the neural signal was significantly different for memorized than for novel trials and thus would corroborate the results of previous studies ^17, 19^.

After averaging the epoched data of correct trials for each experimental condition and combining the planar gradiometers, paired-samples t-tests were performed to identify which condition (memorized or novel) generated a stronger neural signal for each time sample and MEG sensor. Cluster-based MCS were then calculated to correct for multiple comparisons. This was performed independently for both tonal and atonal data (see Methods for details).

First, paired-samples t-tests (_α_ = .01) were calculated for the tonal data in the time interval 0 – 2500 ms (from the onset of the trial) using combined planar gradiometers as these sensors are less affected by external noise than magnetometers ^42-45^. Next, multiple comparisons were corrected by using cluster-based MCS on the significant t-tests’ results (_α_ = .001, 1000 permutations). Three main significant clusters of activity were identified in three specific time intervals when contrasting memorized versus novel sequences, as reported in **Table 1** and in Supplementary Materials (**Figure SF1** and **Table ST1**). Additionally, two main significant clusters of activity were detected when contrasting novel versus memorized sequences (**Table 1, Figure SF2**, and **Table ST1**).

**Table 1.** Significant clusters of activity for the tonal MEG sensor data.

| Cluster number | Size | MEG channels | Time interval (seconds) | *p*-value |
| --- | --- | --- | --- | --- |
| *Memorized versus novel tonal sequences* |  |  |  |  |
| 1 | 224 | 63 | 0.14 – 0.187 | <.001 |
| 2 | 180 | 29 | 0.987 – 1.153 | <.001 |
| 3 | 150 | 37 | 0.807 – 0.887 | <.001 |
| *Novel versus memorized tonal sequences* |  |  |  |  |
| 1 | 277 | 36 | 0.64 – 0.8 | <.001 |
| 2 | 242 | 30 | 0.38 – 0.513 | <.001 |

Regarding the atonal data, paired-samples t-tests (_α_ = .01) were calculated in the same time interval (0 – 2500 ms) using combined planar gradiometers. Next, multiple comparisons were corrected for by using MCS on the significant t-tests’ results (_α_ = .001, 1000 permutations). This procedure identified three main significant clusters of activation when contrasting memorized versus novel excerpts (**Table 2, Figure SF3**, and **Table ST1**). In the case of the novel versus memorized contrast, three main significant clusters of activity were found (**Table 2, Figure SF4**, and **Table ST1**).

**Table 2.** Significant clusters of activity for the tonal MEG sensor data.

| Cluster number | Size | Channels | Time interval (seconds) | *p*-value |
| --- | --- | --- | --- | --- |
| *Memorized versus novel atonal sequences* |  |  |  |  |
| 1 | 288 | 40 | 0.68 – 0.9 | <.001 |
| 2 | 215 | 44 | 0.52 – 0.66 | <.001 |
| 3 | 135 | 40 | 0.133 – 0.187 | <.001 |
| *Novel versus memorized atonal sequences* |  |  |  |  |
| 1 | 478 | 52 | 1.167 – 1.267 | <.001 |
| 2 | 345 | 42 | 0.893 – 0.987 | <.001 |
| 3 | 320 | 44 | 0.653 – 0.74 | <.001 |

### Source reconstruction

After examining the strength of the neural signals at the MEG sensor level, we focused on the main aim of the study, namely to investigate the neural differences underlying the recognition of tonal versus atonal musical sequences in MEG reconstructed source space. To perform this analysis, we localized the brain sources of the neural signal recorded by the MEG channels. This was performed for both the tonal and atonal data and for two frequency bands (delta [0.1 – 1 Hz] and theta [2 – 8 Hz]) that were previously described by Bonetti et al. ^38, 40^ and presumably linked to the processing of the single components (theta) relative to the wholistic sequence (delta).

#### Delta band (0.1 – 1 Hz)

The neural sources were calculated using a beamformer approach. First, a forward model was computed by considering each brain source as an active dipole and calculating its strength across the MEG sensors. Second, a beamforming algorithm was used as an inverse model to estimate the brain sources of the neural activity based on the MEG recordings.

After computing the neural sources, a GLM was calculated at each timepoint and dipole location. A series of t-tests (α = .05) was carried out at the first and group level to estimate the main effect of memorized and novel conditions and their contrast for both the tonal and atonal data independently. Cluster-based MCS (α = .001, 1000 permutations) were computed to correct for multiple comparisons and to determine the brain activity underlying the development of the musical sequences. These analyses were carried out for five specific time intervals that corresponded to each of the tones comprising the sequences: first tone (0 – 250 ms), second tone (251 – 500 ms), third tone (501 – 750 ms), fourth tone (751 – 1000 ms), and fifth tone (1001 – 1250 ms). This was estimated for the memorized versus novel contrast for both tonal and atonal sequences independently and for memorized tonal versus memorized atonal sequences.

Significant clusters of activity (*p* < .001) were located across a number of brain voxels (*k*) for each tone of the tonal sequences, as reported in **Table ST2**. For memorized tonal sequences, the neural activity was overall stronger for the third (*k* = 69), fourth (*k* = 266), and fifth tones (*k* = 229). The largest differences were localized in the middle cingulate gyrus, right supplementary motor area, precuneus, and left lingual gyrus for the third tone; the left amygdala, left parahippocampal gyrus, left lingual gyrus, left hippocampus, and middle cingulate gyrus for the fourth tone, and the anterior and middle cingulate gyrus and left lingual gyrus for the last tone. For novel tonal sequences, the brain activity was stronger for the first (*k* = 54) and second tones (*k* = 29). In particular, the difference between novel and memorized sequences was strongest in the left calcarine fissure, left lingual gyrus, left hippocampus, left precuneus, and left superior temporal gyrus for the first tone, and the right fusiform gyrus, right lingual gyrus, and right inferior occipital gyrus for the second tone. The contrast between memorized and novel tonal sequences for the delta band is depicted in **Figure 3A**.

**Figure 3.**
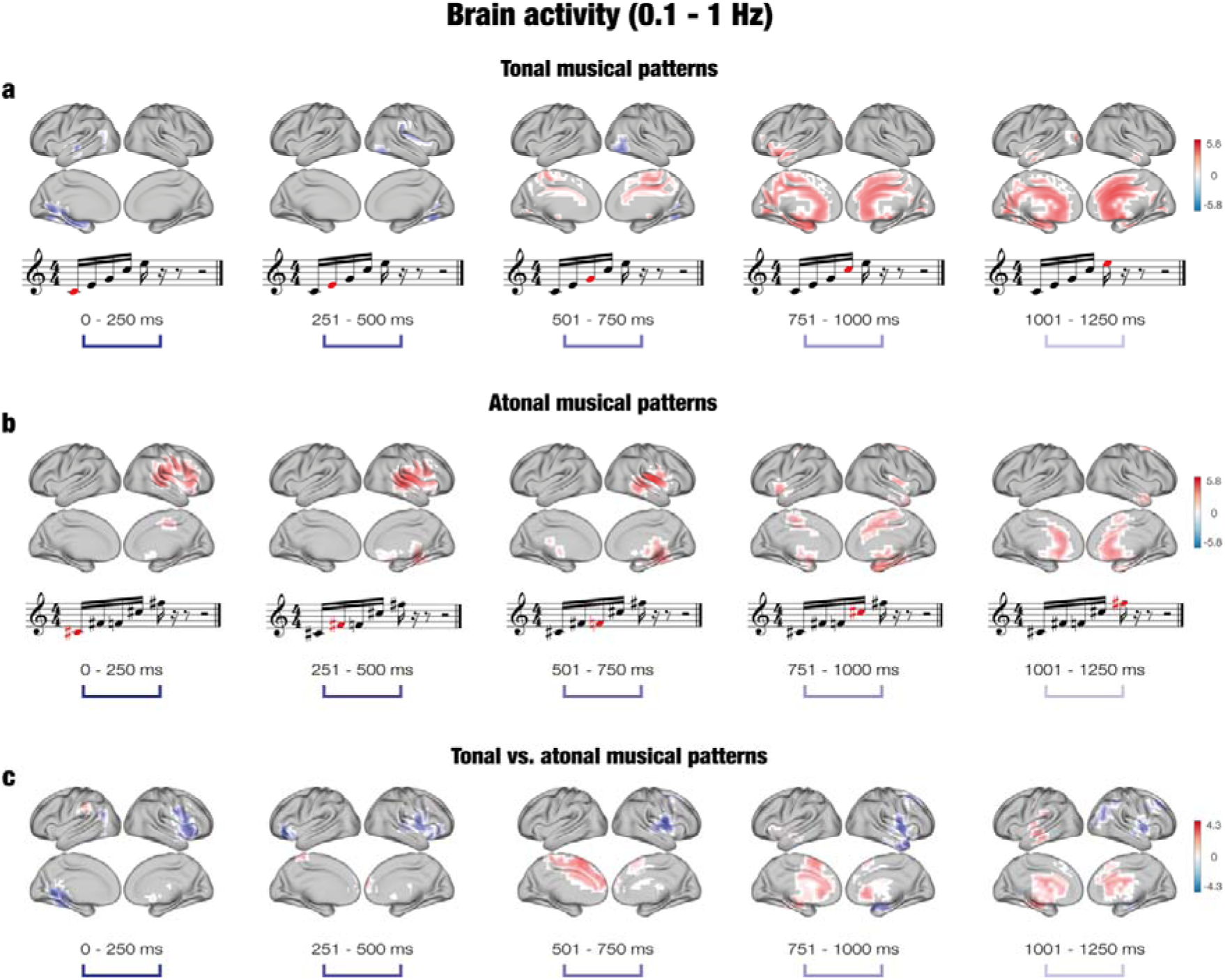
Brain activity underlying the recognition of musical sequences at the delta band (0.1 – 1 Hz) **a** – For tonal sequences, the brain activity was stronger for memorized than novel sequences, particularly for the third (501 – 750 ms), fourth (751 – 1000 ms), and fifth (1001 – 1250) tones. The difference was localized in memory processing areas such as the cingulate gyrus, hippocampus, and parahippocampal gyrus. **b** – For atonal sequences, the brain activity was stronger for memorized than novel sequences for all tones. The difference was mainly localized in auditory processing areas (e.g., superior temporal gyrus, Heschl’s gyrus) for the first three tones, and in memory processing areas (e.g., parahippocampal gyrus, hippocampus) for the fourth and fifth tones. **c** – For the contrast between tonal and atonal sequences, the brain activity was localized in memory processing areas for tonal sequences, particularly for the last three tones, and in auditory processing areas for atonal sequences for all tones. In the case of atonal sequences, significant clusters of activity were located for memorized sequences primarily in the right hemisphere, and the neural activity was stronger for memorized than novel sequences across all five tones (k_1_ = 132, k_2_ = 163, k_3_ = 130, k_4_ = 140, k_5_ = 64), as reported in **Table ST2**. In particular, the brain activity was strongest in the right Rolandic operculum, right superior temporal gyrus, right Heschl’s gyrus, right supramarginal gyrus, and right insula for the first tone; the right Heschl’s gyrus, right superior temporal gyrus, right Rolandic operculum, right middle temporal gyrus, and right insula for the second tone; the right putamen, right insula, right Rolandic operculum, right Heschl’s gyrus, and right thalamus for the third tone; the parahippocampal gyrus, right fusiform gyrus, right hippocampus, and putamen for the fourth tone; and the anterior cingulate cortex, middle frontal gyrus, and caudate nucleus for the last tone. No significant clusters of activity were located in the delta band for novel atonal sequences. **Figure 3B** pictures the contrast between memorized and novel atonal sequences in the delta band.

Regarding the contrast between memorized tonal and atonal sequences, significant clusters of activity were located for both types of musical sequences across all tones (see **Table ST2**). For tonal sequences, the number of significant voxels was higher for the third (*k* = 70) and fifth tones (*k* = 79), whereas for atonal sequences the number of significant brain voxels was higher for the first (*k* = 98), second (*k* = 80), and fourth tones (*k* = 103). In the case of memorized tonal sequences, the neural activity was localized in the the supplementary motor area, left median cingulate gyrus, and superior frontal gyrus for the third tone, and the left hippocampus, left superior temporal gyrus, left thalamus, left insula, left putamen, and left parahippocampal gyrus for the fifth tone. For memorized atonal sequences, the neural activity was localized in the left lingual gyrus, left precuneus, left calcarine fissure, middle temporal gyrus, and right insula at the first tone; the inferior frontal gyrus, right precentral gyrus, right Rolandic operculum, and right superior temporal gyrus for the second tone; the right Rolandic operculum, right middle frontal gyrus, right postcentral gyrus, right putamen, and right insula for the fourth tone; and the right middle frontal gyrus, right angular gyrus, and right thalamus for the last tone. The contrast between memorized tonal and atonal sequences in the delta band is shown in **Figure 3C**.

#### Theta band (2 – 8 Hz)

The same procedure was carried out for assessing the brain activity underlying the recognition of musical sequences in the fast frequency band (2 – 8 Hz). Once the GLM was computed, cluster-based MCS (α = .001, 1000 permutations) were calculated for five time intervals corresponding to each of the five tones that formed the sequence. Again, this was estimated for the memorized versus novel contrast for both tonal and atonal sequences and memorized tonal versus memorized atonal sequences.

Regarding the contrast for tonal sequences, significant clusters of activity (*p* < .001) were located in multiple brain voxels for both memorized and novel sequences, as reported in **Table ST2**. For memorized tonal sequences, the neural activity was overall stronger for the first tone (*k* = 74), whereas it was stronger for novel tonal sequences for the second (*k* = 36), third (*k* = 200), fourth (*k* = 196), and fifth tones (*k* = 70). The brain activity was localized in the right Rolandic operculum, right insula, right Heschl’s gyrus, and right superior temporal gyrus for the first tone for memorized tonal sequences. For novel tonal sequences, the main active areas were the left superior temporal gyrus, insula, Heschl’s gyrus, and left hippocampus for the second tone; Heschl’s gyrus, superior temporal gyrus, insula, and putamen (k = 11) for the third tone; right Heschl’s gyrus, right insula, right Rolandic operculum, and right superior temporal gyrus for the fourth tone; and the right Rolandic operculum, right Heschl’s gyrus, right hippocampus, and right thalamus for the fifth tone. **Figure 4A** displays the contrast between memorized and novel tonal sequences.

**Figure 4.**
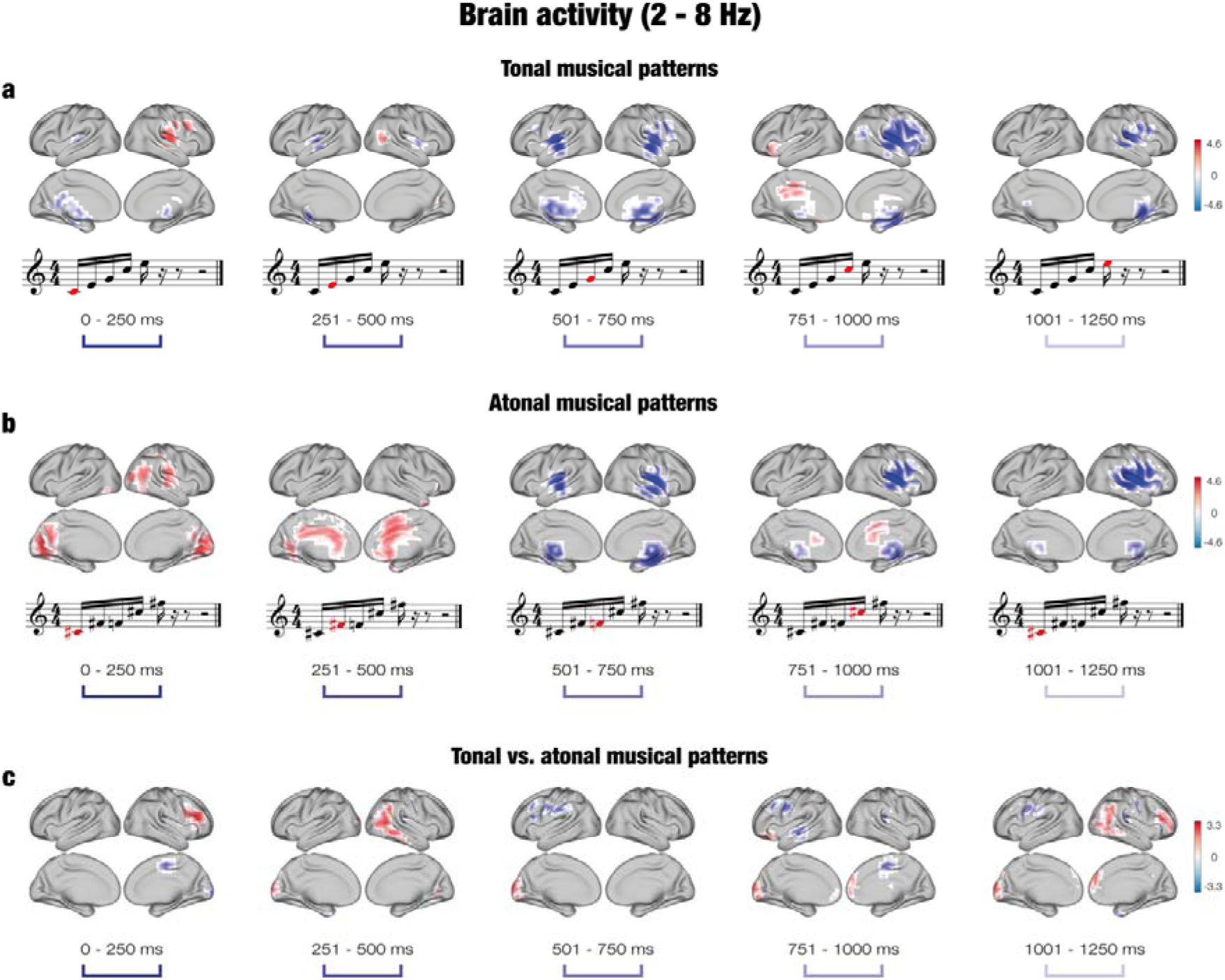
Brain activity underlying the recognition of musical sequences at the theta band (2 – 8 Hz) **a** – For tonal sequences, the brain activity was stronger for novel than memorized sequences, particularly for the last three tones. The difference was localized in auditory processing areas such as Heschl’s gyrus, superior temporal gyrus, and Rolandic operculum. The brain activity was stronger for memorized than novel sequences for the first, second and fourth tones in areas such as the Rolandic operculum (tone 1), occipital gyrus (tone 2) and inferior frontal gyrus (tone 4). **b** – For atonal sequences, the brain activity was stronger for novel than memorized sequences for the last three tones in areas such as the insula, Heschl’s gyrus, and superior temporal gyrus. The brain activity was stronger for memorized than novel sequences for the first two tones in the calcarine fissure and cingulate gyrus. **c** – For the contrast between tonal and atonal sequences, the brain activity was mostly scattered and weak, but the neural activity was stronger in the frontal gyrus (tones 1, 4, and 5), temporal gyrus (tones 2 and 3), and occipital gyrus (tones 2 – 5) for tonal memorized sequences, and in the supplementary motor area (tone 1), frontal gyrus (tones 3 and 4), middle temporal gyrus (tone 4) and postcentral gyrus (tone 5) for atonal memorized sequences.

For the contrast between memorized and novel atonal sequences, the majority of significant clusters of activity were localized for the first (*k* = 166) and second tones (*k* = 104) for memorized sequences, and for the third (*k* = 189), fourth (*k* = 118), and fifth tones (*k* = 156) for novel sequences, as reported in **Table ST2**. For memorized atonal sequences, the neural activity was strongest at the calcarine fissure, lingual gyrus, and right Rolandic operculum for the first tone, and the cingulate gyrus, right supplementary motor area, and superior frontal gyrus for the second tone. For novel atonal sequences, the neural activity was strongest at the insula, putamen, superior temporal gyrus, and Heschl’s gyrus for the third tone; the right insula, right putamen, right Heschl’s gyrus, right Rolandic operculum, and right superior temporal gyrus for the fourth tone; and the right insula, right Rolandic operculum, right Heschl’s gyrus, right putamen, and right superior temporal gyrus for the fifth tone. The contrast between memorized and novel atonal sequences for the theta band is shown in **Figure 4B**.

Finally, the significant clusters of activity in the tonal versus atonal contrast for the theta band are reported in **Table ST2**. In the case of memorized tonal sequences, the number of significant brian voxels was higher for the first (*k* = 20), second (*k* = 44), fourth (*k* = 45), and fifth tones (*k* = 71). The neural activity was located in the right inferior frontal gyrus and right middle frontal gyrus for the first tone; the right middle temporal gyrus, right inferior parietal gyrus, right angular gyrus, middle occipital gyrus, and left superior occipital gyrus for the second tone; the frontal gyrus for the fourth tone; and the right middle occipital gyrus, right frontal gyrus, and right middle temporal gyrus for the fifth tone. For memorized atonal sequences, the number of significant voxels was higher for the third tone (*k* = 38) and the neural activity was mainly localized in the left inferior frontal gyrus, left middle frontal gyrus, left supramarginal gyrus, and right supplementary motor area. **Figure 4C** shows the contrast between tonal and atonal sequences for the theta band.

Altogether, we found significant differences between tonal and atonal sequences, especially for the slow frequency band. Recognition of memorized tonal sequences elicited stronger neural activity in left cingulate and hippocampal areas in the last three tones of the sequences, whereas recognition of memorized atonal sequences was supported by activation in right auditory regions from the second tone onwards.

## Discussion

This study set out to investigate the brain activation underlying the recognition of auditory musical sequences characterized by different levels of complexity (tonal and atonal). Behavioral data showed clear differences between the recognition of tonal and atonal sequences and significant clusters of activation were observed at the MEG sensor level. Source reconstruction analyses indicated different activation clusters for tonal and atonal sequences, particularly in the delta frequency band. Overall, the neural activity was stronger in memory processing areas for memorized tonal sequences and in auditory processing regions for memorized atonal sequences.

Prior to focusing on the differences in brain activity related to distinct levels of recognition complexity, we verified that the current results were consistent with previous studies. Indeed, the brain areas activated during the recognition of tonal sequences confirmed the involvement of a widespread brain network including both auditory and memory processing regions ^36-38, 40^. Furthermore, in accordance with previous research, the neural activity was clearly distributed in two frequency bands ^38, 39^. Delta band (0.1 – 1 Hz) was linked to the recognistion of the whole auditory sequences (*global processing*), which was reflected by the stronger activation occurring in this band for the recognition of the memorized sequences. Conversely, theta band (2 – 8 Hz) was associated with the processing of the individual tones (*local processing*), as suggested by the stronger neural activity in auditory regions during processing of novel sequences.

Regarding the recognition of tonal and atonal sequences, we observed distinct neural pathways when processing and recognizing the two types of auditory stimuli. While the recognition of tonal sequences mainly recruited a widespread brain network involving cingulate gyrus and hippocampus in the right hemisphere, the recognition of atonal sequences was mainly associated with a sustained, slow activity in the left auditory cortex. These results can be interpreted in light of different theoretical frameworks, namely predictive coding, harmonicity, and global neuronal workspace (GNW). According to PCM theory, the brain’s predictive model is being continuously updated while listening to music in order to decrease precision-weighted prediction errors ^16, 18, 19^. The predictive value of atonal music is weaker than tonal music, which alters its complexity and increases prediction errors ^42-46, 49^. In turn, this change in stimulus predictability undermines the processing ^42-46^ and enjoyment ^46-48^ of atonal music. This was apparent when examining the behavioral results, since memorized atonal sequences were more slowly and less accurately recognized. In addition, the distribution of the neural activity in two frequency bands suggests a combination of top-down predictions in the delta band, which is related to the recognition of memorized sequences, and bottom-up predictions in the theta band, as the prediction error increases with novel sequences.

An alternative explanation for these results focuses on the harmonicity of auditory stimuli. Tonal music has been closely linked to the harmonic series, a natural sequence of sound frequencies that are integer multiples of a fundamental. Environmental sounds are typically nonharmonic, whereas both human and animal vocalizations contain harmonic structures ^63^. The tonotopic organization of the human auditory cortex is particularly sensitive to harmonic tones, suggesting that this region developed to process harmonics due to their relevance for social communication ^64, 65^. These results indicate that distinct neural pathways are activated when recognizing auditory stimuli that are not coherent with the natural harmonic series and thus arguably more complex to process. Indeed, we found that for memorized tonal sequences, the brain activity was primarily located at the cortico-hippocampal network in the right hemisphere, and for memorized atonal sequences the auditory network in the left hemisphere. Specifically, the strong activation in low-processing primary auditory regions at the first three tones of the atonal sequences suggests a “disentangling” of the sequence before it can be processed and recognized by high-cognitive areas involved in memory processing. One possible approach to further test the harmonicity hypothesis in relation to sequence recognition would be to create a collection of pieces that are systematically varied in terms of their similarity to the natural harmonic series. Future studies are called to investigate such perspectives.

Finally, the current results are also consistent with the GNW hypothesis ^66, 67^. According to this theory, stimuli become conscious when they ignite late, high-order regions in response to the activation of sensory cortices involved in perceptual representation. Conversely, unconscious information does not reach high-processing brain areas and neural activity is limited to sensory cortices ^66, 68, 69^. Importantly, we found that tonal sequences induced a late and robust activation of memory processing regions. Although it is unclear *why* atonal sequences were differently processed by the brain, we can confirm that the complexity of the stimuli modulates the transition from primary sensory areas to the GNW, adding new information to this comprehensive theoretical framework.

The current study provides valuable insights into the brain mechanisms underlying the recognition of auditory sequences. The results are consistent with those of previous studies and evidence of the engagement of a large brain network that comprises both memory processing and auditory regions when recognizing music. Results further highlight the importance of stimulus complexity for the processing of temporal sequences and hint that the brain employs different strategies to account for this complexity.

## Supporting information

This file includes four supplementary and their descriptions, along with the descriptions for the two supplementary tables.

This table reports the significant clusters of activity for the magnetoencephalography sensor data.

This table reports the significant clusters of activity for the magnetoencephalography source data.

## Acknowledgements

We thank Giulia Donati, Riccardo Proietti, Giulio Carraturo, Mick Holt, Holger Friis for their assistance in the neuroscientific experiment.

The Center for Music in the Brain (MIB) is funded by the Danish National Research Foundation (project number DNRF117). Additionally, we thank the Fundación Mutua Madrileña for the economic support provided to the author GFR and the University of Bologna for the economic support provided to student assistants Giulia Donati, Riccardo Proietti and Giulio Carraturo.

LB is supported by Carlsberg Foundation (CF20-0239), Center for Music in the Brain, Linacre College of the University of Oxford, and Society for Education and Music Psychology (SEMPRE’s 50th Anniversary Awards Scheme).

MLK is supported by Center for Music in the Brain and Centre for Eudaimonia and Human Flourishing, which is funded by the Pettit and Carlsberg Foundations.

## Author contributions

LB, EB, MLK, and PV conceived the hypotheses, designed the study, and recruited the resources for the experiment. LB and GFR performed pre-processing and statistical analysis. EB, SAK, LB, MLK, and PV provided essential help to interpret and frame the results within the neuroscientific literature. GFR and LB wrote the first draft of the manuscript and prepared the figures. All the authors contributed to and approved the final version of the manuscript.

## Materials and methods

### Participants

The participant sample consisted of 71 volunteers (38 males and 33 females) aged 18 to 42 years old (mean age: 25 ± 4.10 years). All participants were healthy and reported normal hearing. Participants were recruited in Denmark and came from Western countries with matching socioeconomic and educational backgrounds.

The project was approved by the Ethics Committee of the Central Denmark Region (De Videnskabsetiske Komitéer for Region Midtjylland, Ref 1-10-72-411-17). The experimental procedures were carried out in compliance with the Declaration of Helsinki – Ethical Principles for Medical Research. All participants gave the informed consent before starting the experimental procedure.

### Experimental stimuli and design

Two musical compositions were used in the experiment: the right-hand part of Johann Sebastian Bach’s Prelude No. 1 in C Major, BWV 846 (hereafter referred to as the “tonal piece”), and an atonal version of the prelude (hereafter referred to as the “atonal piece”). MIDI versions were created using Finale (MakeMusic, Boulder, CO) and both pieces lasted 2.5 minutes each, with the same duration for all tones. LB composed the atonal piece based on the tonal piece. In particular, new tones were assigned to each of the tones comprising Bach’s original prelude. These new tones were one or two semitones higher or lower than the original tones, and the same tone conversion was applied throughout the entire tonal piece to obtain the atonal piece (e.g., every C tone in the tonal piece was converted into a C sharp in the atonal piece). Thus, both compositions were identical in terms of the sequential presentation of the tones (i.e., if C was positioned as 1^st^, 7^th^, and 8^th^ tone in the tonal piece, C sharp occupied the same positions [1^st^, 7^th^, and 8^th^] in the atonal piece), their rhythmic pattern, dynamics, and duration. Thus, the crucial difference between the two pieces was that the tonal piece was in the key of C Major, whereas the atonal piece did not have a musical key. The first two bars of each piece are displayed in **Figure 1a**, showing similarities and correspondence between the two pieces.

Forty musical excerpts (i.e., short melodies or sequences) were extracted from each of the pieces. All excerpts consisted of the first five notes of each bar and lasted for 1250 ms (250 ms per note). In addition, 40 new excerpts were created for each piece based on the original ones. These new sequences were matched to the original ones among several variables, to prevent potential confounds. Specifically, they were matched for rhythm, volume, timbre, tempo, meter, and tonality.

The stimuli were employed in an old/new auditory recognition paradigm, as depicted in **Figure 1b**, that was administered to the participants while their brain activity was recorded using MEG. The paradigm consisted of two parts, encoding and recognition, and was performed twice, once for the tonal piece and once for the atonal piece. The order of tonal/atonal was counterbalanced across participants. During the encoding part, participants actively listened to four repetitions of the entire musical piece (tonal or atonal) and tried to memorize it as much as possible. Afterwards, they were presented with the previously described 80 musical excerpts (40 memorized and 40 novel excerpts, randomly ordered) and stated whether the excerpts belonged to the piece they had previously listened to (“memorized”) or whether they were new excerpts (“novel”). Response accuracy and reaction time were recorded using a joystick.

### Data acquisition

The MEG recordings were acquired in a magnetically shielded room at Aarhus University Hospital (Denmark) with an Elekta Neuromag TRIUX MEG scanner with 306 channels (Elekta Neuromag, Helsinki, Finland). The data were recorded at a sampling rate of 1000 Hz with an analogue filtering of 0.1 – 330 Hz. Before starting the recordings, the head shape of the participants and the position of four Head Position Indicator (HPI) coils with respect to three anatomical landmarks were registered using a 3D digitizer (Polhemus Fastrak, Colchester, VT, USA). This information was later used to co-register the MEG data with the MRI anatomical scans. During the MEG recordings, the HPI coils registered the continuous head localization, which was subsequently used for movement correction analyses. Additionally, two sets of bipolar electrodes were used to record eye movements and cardiac rhythm for later removing electrooculography (EOG) and electrocardiography (ECG) artifacts.

The MRI scans were recorded on a CE-approved 3T Siemens MRI-scanner at Aarhus University Hospital (Denmark). The data were recorded using a structural T1 with a spatial resolution of 1.0 × 1.0 × 1.0 mm and the following sequence parameters: echo time (TE) = 2.96 ms, repetition time (TR) = 5000 ms, reconstructed matrix size = 256 × 256, bandwidth = 240 Hz/Px.

The MEG and MRI recordings were acquired in two separate sessions.

### Data preprocessing

The raw MEG sensor data (204 planar gradiometers and 102 magnetometers) were first preprocessed by MaxFilter ^50^ in order to suppress external interferences. In addition, the data were corrected for head motion and downsampled to 250 Hz. The data were then converted into Statistical Parametric Mapping (SPM) ^51^ format and analyzed in MATLAB (MathWorks, Natick, MA, USA) with the Oxford Centre for Human Brain Activity (OHBA) Software Library (OSL) (https://ohba-analysis.github.io/osl-docs/), a freely available software that builds upon Fieldtrip ^52^, FSL ^53^, and SPM toolboxes. The signal was high-pass filtered (0.1 Hz of cutoff) to remove external frequencies and a notch filter was subsequently applied (48 – 52 Hz) to correct for inferences of the electric current. The signal was further downsampled to 150 Hz and the continuous MEG data were visually inspected to remove artifacts using the OSLview tool. An independent component analysis (ICA) was performed to remove EOG and ECG components. After reconstructing the signal with the remaining components ^54^, the data were epoched into 160 trials (80 excerpts from each musical piece). Each trial lasted 1350 ms (1250 ms plus 100 ms of baseline time) and further analyses were performed on correctly identified trials only (see **Figure 1C**).

The raw MEG sensor data (204 planar gradiometers and 102 magnetometers) were first preprocessed by MaxFilter ^50^ in order to suppress external interferences. In addition, the data were corrected for head motion and downsampled to 250 Hz. The data were then converted into Statistical Parametric Mapping (SPM) ^51^ format and analyzed in MATLAB (MathWorks, Natick, MA, USA) with the Oxford Centre for Human Brain Activity (OHBA) Software Library (OSL) (https://ohba-analysis.github.io/osl-docs/), a freely available software that builds upon Fieldtrip ^52^, FSL ^53^, and SPM toolboxes. The signal was high-pass filtered (0.1 Hz of cutoff) to remove external frequencies and a notch filter was subsequently applied (48 – 52 Hz) to correct for inferences of the electric current. The signal was further downsampled to 150 Hz and the continuous MEG data were visually inspected to remove artifacts using the OSLview tool. An independent component analysis (ICA) was performed to remove EOG and ECG components. After reconstructing the signal with the remaining components ^54^, the data were epoched into 160 trials (80 excerpts from each musical piece). Each trial lasted 1350 ms (1250 ms plus 100 ms of baseline time) and further analyses were performed on correctly identified trials only (see **Figure 1C**).

### MEG sensor data analysis

The primary focus of this study was on detecting differences in the brain activity underlying the recognition of tonal versus atonal musical sequences. However, the data were first analyzed at the MEG sensor level to verify that the neural signal was stronger for memorized versus novel musical sequences. This first step was essential to replicate previous findings obtained using a very similar experimental setting and paradigm and thus assess the quality of our data ^38-40^.

Following the preprocessing of the neural data, and in accordance with MEG analysis guidelines ^55^, all trials belonging to one condition were averaged together. This procedure resulted in four mean trials: one for memorized trials and one for novel trials for each musical piece (i.e., memorized tonal, novel tonal, memorized atonal, novel atonal). Next, each pair of planar gradiometers was combined by a sum root square. Paired-samples t-tests (α = .01) were then calculated to contrast the memorized and novel conditions for both the tonal and atonal pieces, independently. This was performed for each combined planar gradiometer and each time-point in the time-range 0 – 2500 ms (from the onset of the first tone of the musical sequences) in order to determine which condition generated a stronger neural signal. The analyses were calculated for planar gradiometers, since these sensors are less affected by external noise and thus highly reliable when computing analyses at the MEG sensor level ^55-58^. Multiple comparisons were corrected using cluster-based Monte Carlo simulations (MCS) ^59^ (α = .001, 1000 permutations) on the significant t-tests’ results. Specifically, for each timepoint, a 2D matrix was generated reproducing the spatial location of the MEG channels and the results of the t-tests of each MEG channel binarized according to their *p*-values (0s for not significant tests and 1s for significant tests [i.e., *p* < .01]). The elements of the resulting 3D matrix were then submitted to 1000 permutations. For each permutation, we identified the maximum cluster of permuted 1s, and we built a reference distribution using the maximum cluster sizes detected for each of the 1000 permutations. Finally, the original clusters that had a larger size than 99.9% of the maximum cluster sizes of the permuted data were considered significant.

### Source reconstruction

After examining the strength of the neural signals at the MEG sensor level, we focused on the main aim of the study, which was to investigate the neural differences underlying the recognition of tonal versus atonal musical sequences in MEG reconstructed source space. To perform this analysis, we localized the brain sources of the neural signal recorded in the MEG channels. This procedure required designing a forward model, computing the inverse solution (in this case, using a beamforming approach), and identifying the statistically significant brain sources underlying the recognition of tonal and atonal sequences and their contrasts over time. **Figure 1D** shows the graphical depiction of the source reconstruction analyses.

#### Beamforming

Before computing the source reconstruction algorithm, the continuous data were band-pass filtered into two frequency bands: a slow band (delta, 0.1 – 1 Hz) and a fast band (theta, 2 – 8 Hz). These bands were selected based on the findings reported by Bonetti et al. ^38-40^, which suggested that the theta band was responsible for a sensorial elaboration of each object (tone) of the sequence, while the delta band was implicated in the recognition of the holistic temporal sequence. The filtered data were then epoched and the brain sources that generated the signal were calculated.

First, an overlapping-spheres forward model was computed using an 8-mm grid. This theoretical head model considers each brain source as an active dipole and describes how the unitary strength of such a dipole is reflected across the MEG sensors ^60^. Using the information collected with the 3D digitizer, the MEG data and individual T1-weighted images were co-registered and the forward model was subsequently computed. An MNI152-T1 template with 8-mm spatial resolution was used in four cases in which the individual anatomical scans were not available. Second, a beamforming algorithm was employed as the inverse model. This is one of the most widely used algorithms for estimating the brain sources from MEG channels’ data and consists of utilizing a different set of weights which are sequentially applied to the source locations (dipoles) for isolating the contribution of each source to the activity recorded by the MEG channels for each time-point ^55, 61, 62^.

#### General Linear Model

After estimating the brain sources of the signal recorded on the MEG channels, a General Linear Model (GLM) was estimated sequentially for each timepoint at each dipole location. At the first level, the main effect of memorized and novel conditions, as well as their contrast, was computed independently for each participant. At the group level, t-tests were carried out for each dipole location to obtain the main effect of tonal, atonal and their contrast computed on all aggregated participants. The GLMs were estimated independently for both the tonal and atonal data and for both frequency bands.

#### Brain activity underlying the development of the musical sequences

To determine the temporal evolution of the brain activity underlying musical sequences’ recognition, cluster-based MCS were estimated for five specific time-windows that corresponded to each of the five tones comprising the musical sequences. This procedure was carried out independently for both tonal and atonal data and for both frequency bands. Thus, ten cluster-based MCS were calculated for each musical piece (five tones x two frequency bands) on the results of the group-level analysis with an adjusted alpha level of .001 (α = 0.01/10 = .001). This procedure allowed detecting the spatial clusters of significant brain sources underlying the recognition of the tonal and atonal musical sequences. For each of the MCSs, the data were sub-averaged in the time-window of interest (e.g., the time-window for the first tone of the musical sequences) and then submitted to 1000 permutations to build a reference distribution of the maximum cluster sizes detected in the permuted data. Then, using the same procedure as with the MEG channels, the original cluster sizes were compared to the reference distribution and were considered significant if their size was bigger than 99.9% of the maximum cluster sizes of the permuted data.

Importantly, further analyses were conducted to assess the differences between tonal and atonal data when recognizing memorized trials for both the delta and theta frequency bands. For each participant, a t-test (α = 0.01) was computed for each source location and for the five time-windows corresponding to each musical tone, contrasting the brain activity underlying the recognition of tonal versus atonal music. Multiple comparisons were corrected for by using cluster-based MCS, as described above. In this case, ten MCS (α = .001, 1000 permutations) were calculated on the significant t-test results (five tones x two frequency ranges).

## Data availability

The code and anonymized neuroimaging data from the experiment will be made available upon request. Regarding the data, we will be able to share it when it is completely anonymized and cannot lead in any way to the original participants identity, according to Danish regulations. Otherwise, a data sharing agreement must be made.

