## Supplementary material for "The spatiotemporal dynamics of recognition memory for complex versus simple auditory sequences": This file includes four supplementary and their descriptions, along with the descriptions for the two supplementary tables.

### *Supplementary materials*

Supplementary materials related to this study are organized as supplementary figures and tables. Due to their large size, supplementary tables have been reported in Excel files.

#### *Supplementary figures*

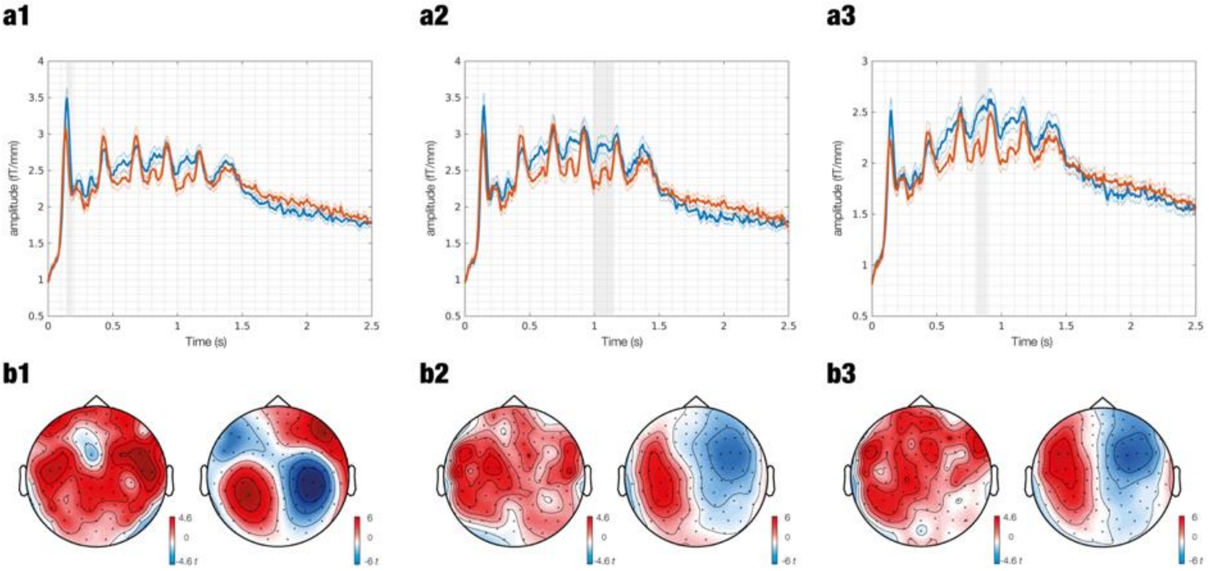


***Figure SF1. Graphical depiction of the significant clusters of activity for tonal MEG sensor data (memorized vs novel musical excerpts)***

**a** – The plots represent the signal amplitude of the tonal memorized patterns (in red) and tonal novel patterns (in blue) in clusters 1 (**a1**), 2 (**a2**), and 3 (**a3**). The plots show the full time-window of analyses (0 to 2.5 seconds). Significant time intervals are marked in grey. **b** – The topoplots show the contrast between tonal memorized patterns (in red) and tonal novel patterns (in blue) for each of the significant time intervals of cluster 1 (**b1**; 0.14 – 0.187 seconds), 2 (**b2**; 0.987 – 1.153 seconds), and 3 (**b3**; 0.807 – 0.887 seconds). The left topoplots depict the neural activity recorded by gradiometeres and the right topoplots show the neural activity recorded by magnetometers.

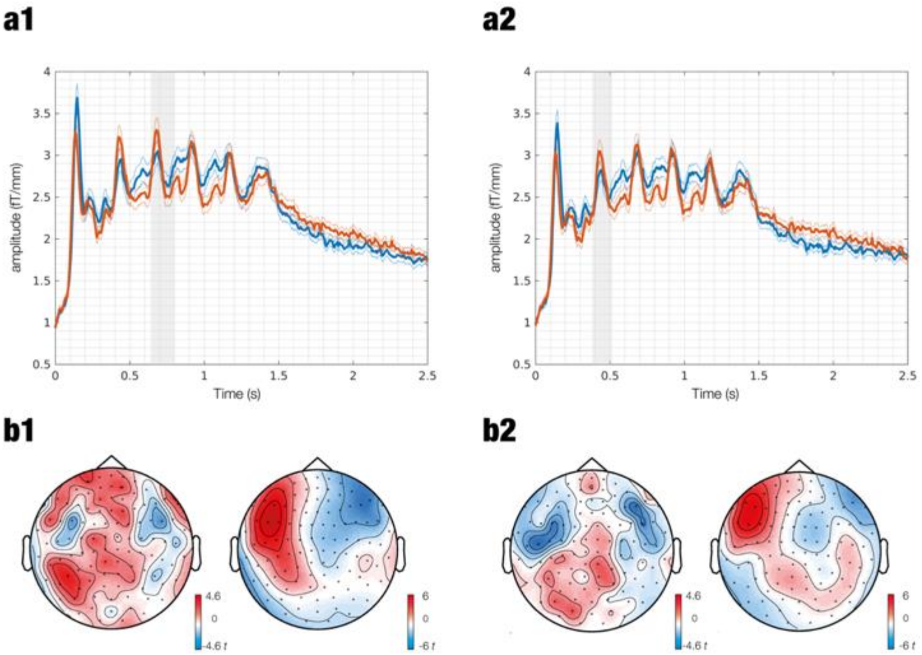


***Figure SF2. Graphical depiction of the significant clusters of activity for tonal MEG sensor data (novel vs memorized musical excerpts)***

**a** – The plots represent the signal amplitude of the tonal memorized patterns (in red) and tonal novel patterns (in blue) in clusters 1 (**a1**) and 2 (**a2**). The plots show the full time-window of analyses (0 to 2.5 seconds). Significant time intervals are marked in grey. **b** – The topoplots show the contrast between tonal novel patterns (in red) and tonal memorized patterns (in blue) for each of the significant time intervals of cluster 1 (**b1**; 0.64 – 0.8 seconds) and 2 (**b2**; 0.38 – 0.513 seconds). The left topoplots depict the neural activity recorded by gradiometeres and the right topoplots show the neural activity recorded by magnetometers.

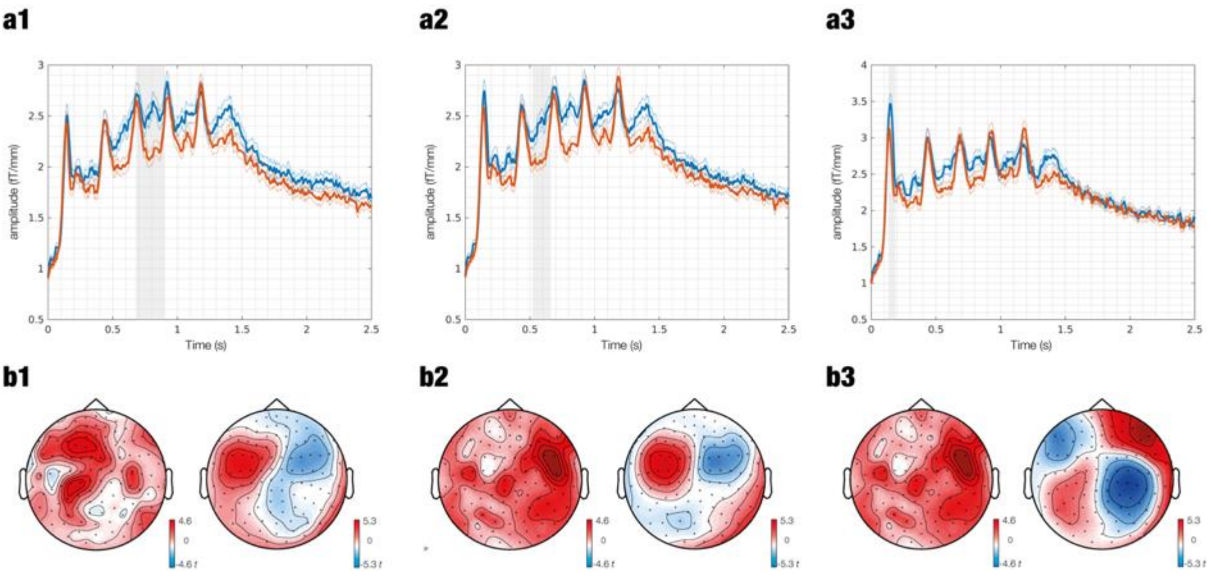


***Figure SF3. Graphical depiction of the significant clusters of activity for atonal MEG sensor data (memorized vs novel musical excerpts)***

**a** – The plots represent the signal amplitude of the atonal memorized patterns (in red) and atonal novel patterns (in blue) in clusters 1 (**a1**), 2 (**a2**), and 3 (**a3**). The plots show the full time-window of analyses (0 to 2.5 seconds). Significant time intervals are marked in grey. **b** – The topoplots show the contrast between atonal memorized patterns (in red) and atonal novel patterns (in blue) for each of the significant time intervals of cluster 1 (**b1**; 0.68 – 0.9 seconds), 2 (**b2**; 0.52 – 0.66 seconds), and 3 (**b3**; 0.133 – 0.187 seconds). The left topoplots depict the neural activity recorded by gradiometeres and the right topoplots show the neural activity recorded by magnetometers.

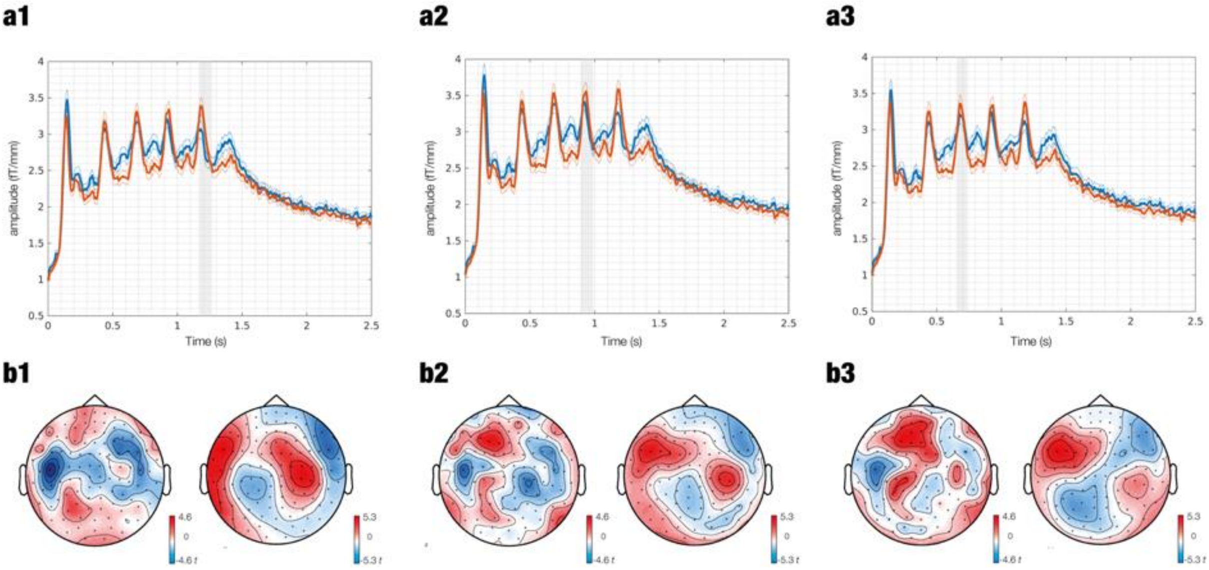


***Figure SF4. Graphical depiction of the significant clusters of activity for atonal MEG sensor data (novel vs memorized musical excerpts)***

**a** – The plots represent the signal amplitude of the atonal memorized patterns (in red) and atonal novel patterns (in blue) in clusters 1 (**a1**), 2 (**a2**), and 3 (**a3**). The plots show the full time-window of analyses (0 to 2.5 seconds). Significant time intervals are marked in grey. **b** – The topoplots show the contrast between atonal memorized patterns (in red) and atonal novel patterns (in blue) for each of the significant time intervals of cluster 1 (**b1**; 1.167 – 1.267 seconds), 2 (**b2**; 0.893 – 0.987 seconds), and 3 (**b3**; 0.653 – 0.74 seconds). The left topoplots depict the neural activity recorded by gradiometeres and the right topoplots show the neural activity recorded by magnetometers.

*Supplementary tables*

***Table ST1. Significant clusters of activity for MEG sensor data***

Significant clusters of activity estimated from the two-sided contrasts between memorized and novel musical patterns. This was performed for both tonal and atonal patterns independently. The Excel file depicts the number of significant clusters, along with the MEG channels and time-windows.

***Table ST2. Significant clusters of activity for MEG source data***

Significant clusters of activity estimated from the contrasts between tonal and atonal memorized musical patterns. This was performed for both delta and theta frequency bands independently. The Excel file depicts the contrast for each of the tones comprising the musical patterns, along with the brain regions, hemispheres, and averaged *t*-values for each voxel.
